# Spontaneous nucleation and fast aggregate-dependent proliferation of α-synuclein aggregates within liquid condensates at physiological pH

**DOI:** 10.1101/2021.09.26.461836

**Authors:** Samuel T. Dada, Maarten C. Hardenberg, Lena K. Mrugalla, Mollie O. McKeon, Ewa Klimont, Thomas C. T. Michaels, Michele Vendruscolo

## Abstract

It is well-established that α-synuclein aggregation may proceed through an initial lipid-dependent aggregate formation and, if at acidic pH, a subsequent aggregate-dependent proliferation. It has also been recently reported that the aggregation of α-synuclein may also take place through an alternative pathway, which takes place within dense liquid condensates produced through liquid-liquid phase separation. The microscopic mechanism of this process, however, remains to be clarified. Here, we developed a fluorescence-based assay to perform a kinetic analysis of the aggregation process of α-synuclein within liquid condensates, and applied it to determine the corresponding mechanism of aggregation. Our analysis shows that at pH 7.4 the aggregation process of α-synuclein within dense condensates starts with spontaneous primary nucleation followed by rapid aggregate-dependent proliferation. Taken together, these results reveal a highly efficient pathway for the appearance and proliferation of α-synuclein aggregates at physiological pH.

## Introduction

Parkinson’s disease is the most common neurodegenerative movement disorder^1,2^. A distinctive pathophysiological signature of this disease is the presence of abnormal intraneuronal protein deposits known as Lewy bodies^3,4^. One of the main components of Lewy bodies is α-synuclein^5^, a peripheral membrane protein highly abundant at neuronal synapses^6,7^, and genetically linked with Parkinson’s disease^8,9^. This 140-residue disordered protein can be subdivided into three domains, an amphipathic N-terminal region (amino acids 1-60), a central hydrophobic region (non-amyloid-β component, or NAC, amino acids 61-95) and an acidic proline-rich C-terminal tail (amino acids 96-140)^7^. Although α-synuclein is typically found in misfolded and aggregated species in Lewy bodies, the corresponding mechanism of its aggregation and role in Parkinson’s disease are not yet fully understood.

Quite generally, the aggregation process of proteins proceeds through a series of microscopic steps, including primary nucleation, elongation, and secondary nucleation^10,11^. During primary nucleation, the self-assembly of proteins in their monomeric form leads to the formation of aggregate seeds, an event that may occur in solution or on lipid surfaces^12,13^. The formation of these seeds is an intrinsically slow event at the microscopic level, which is observable macroscopically as a lag phase in the aggregation process^10,11^. These initially disordered seeds may then dissociate or convert into ordered assemblies rich in β structure^14^. This conversion step is followed by a growth phase in which the ordered seeds elongate into protofibrils and eventually insoluble fibrillar aggregates. The surfaces of existing fibrillar aggregates then further catalyse the formation of more aggregate seeds^15,16^. This process, which is called secondary nucleation, is typically characterised by the assembly of protein monomers on the surface of fibrils that eventually nucleate into new oligomeric species^15,16^. Once a critical concentration of fibrils is reached, a positive feedback mechanism allows for a rapid cascade of fibril proliferation by autocatalytic processes^15^.

In the case of the aggregation process of α-synuclein, several questions are still open, including two that we are going to address in this study. The first concerns whether there are cellular conditions under which α-synuclein can undergo spontaneous aggregation, and the second whether the proliferation of α-synuclein fibrils by aggregate-dependent feedback processes can take place at physiological pH. According to our current knowledge, α-synuclein aggregation under physiological conditions does not readily take place spontaneously, but it can be initiated by lipid membranes. Furthermore, secondary nucleation contributes significantly to the aggregation process only at acidic pH^13,17^. At this level of understanding, it remains challenging to rationalise the process by which α-synuclein aggregation is linked with Parkinson’s disease.

In order to make further progress, in this work we investigated whether it is possible to leverage the recent finding that α-synuclein can undergo a liquid-liquid phase separation process resulting in the formation of dense liquid condensates^18–20^. Liquid-liquid phase separation has recently emerged as a general phenomenon associated with a wide variety of cellular functions^21–24^ and closely linked with human disease^22,25–27^. A liquid-liquid phase separation process has been reported for a wide range of proteins implicated neurodegenerative conditions, including tau, FUS and TDP-43^28–30^. Since it has also been shown that protein aggregation can take place within liquid condensates (or droplets)^19,25,30–34^, we asked whether it is possible to characterise at the microscopic level the droplet-induced aggregation mechanism of α-synuclein.

To enable the accurate determination of the rate constants for α-synuclein primary and secondary nucleation within droplets, we developed a strategy based a fluorescence-based aggregation assay to monitor both the spontaneous aggregation of α-synuclein and the aggregation in the presence of aggregate seeds. Using this assay within the framework of a kinetic theory of protein aggregation^10,11,35^, we show that α-synuclein can undergo spontaneous primary nucleation and fast aggregate-dependent aggregate proliferation within droplets at physiological pH.

## Results and Discussion

### Prerequisites for the study of α-synuclein aggregation following liquid-liquid phase separation

Increasing evidence indicates that aberrant protein aggregation can take place within dense liquid condensates generated through liquid-liquid phase separation^19,25,30–33^. Fluorescence recovery after photobleaching (FRAP) can be employed to monitor the progressive maturation of the liquid condensates, a process known as Ostwald ripening^36^, and the subsequent appearance of solid-like assemblies^18,19^ (**Supplementary Figures 1** and **2**, and **Supplementary Movies 1** and **2**). The accurate time-dependent measurement of the formation of ordered aggregate species, however, is still challenging. Therefore, the study of α-synuclein aggregation within condensates requires the development of a comprehensive strategy to both captures the initial stages of liquid-liquid phase separation, the maturation of the resulting condensates, as well as the key microscopic steps that drive the subsequent conversion into amyloid aggregates.

### An in vitro strategy for the study of α-synuclein aggregation within liquid condensates

To monitor closely the microscopic steps that drive α-synuclein aggregation within liquid condensates, we built on an established assay used to observe rapid liquid-liquid phase separation of α-synuclein *in vitro^19^*. This assay enables the monitoring of α-synuclein condensates over time and is conducted by depositing a small sample volume as a droplet on a microscopy slide, where the phase separation process is initiated at physiological pH through the use of a molecular crowding agent, in our case PEG (**Figure 1**). Liquid-liquid phase separation of α-synuclein can subsequently be characterised in real-time with the use of fluorescence microscopy using a confocal microscope and α-synuclein labeled with Alexa Fluor 647. Next, to obtain a robust readout of the conversion of α-synuclein from its monomeric state to the amyloid state, we implemented the use of thioflavin T (ThT), a well-known benzothiazole fluorescent amyloid-binding dye that has been employed for both *in vitro* and *in vivo* characterisation of amyloid formation; ThT exhibits an enhanced fluorescent signal upon binding cross-β-strand structures, such as those found in amyloid fibrils^10,19^. Our results indicate that ThT can respond to changes in concentration of preformed α-synuclein fibrils in a concentration-dependent manner and under the specific experimental conditions of our liquid-liquid phase separation assay (**Supplementary Figure 3**).

**Figure 1.**
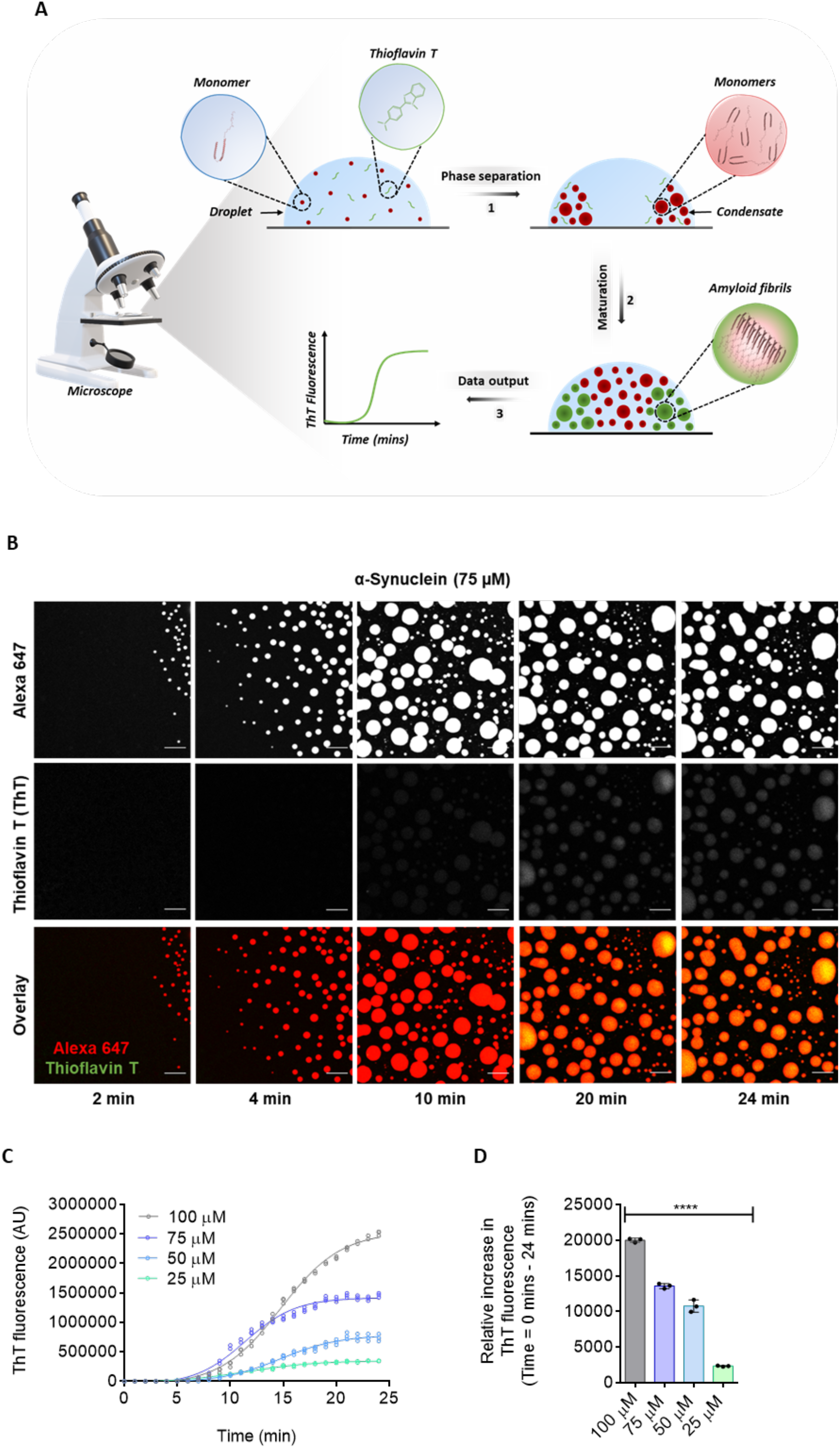
Development of a ThT-based assay to monitor α-synuclein aggregation within liquid condensates. **(A)** The assay has three components: **(1)** the phase separation of α-synuclein into dense liquid droplets (condensates) within dilute liquid droplets is monitored using Alexa Fluor 647 fluorescence, (2) upon maturation of the dense liquid droplets, α-synuclein forms amyloid assemblies over time, and (3) the real-time assessment of amyloid formation is performed using ThT fluorescence. **(B)** Imaging of α-synuclein condensate formation (using Alexa Fluor 647 fluorescence) and aggregation (using ThT fluorescence) over time. In the presence of a crowding agent (10% PEG), α-synuclein monomers (75 μM) form a dense liquid phase surrounded by a dilute liquid phase. In the presence of 20 μM ThT, amyloid-containing condensates can be detected as an increase of the ThT fluorescent signal over time. Scale bar represents 10 μm. **(C)** Quantification of ThT fluorescence over time of the images shown in panel B for 75 μM (blue) α-synuclein. Subsequently, ThT emission for 100 μM (grey), 50 μM (cyan) and 25 μM (turquoise) α-synuclein were obtained in the same manner over a 24-min time period. The curve represents a non-linear fit of the ThT signal (integrated density) over time. **(D)** Relative increase in the raw data values of the ThT fluorescence intensities from 0 to 24 min. All experiments were performed in 50 mM Tris-HCl (pH 7.4) in the presence of 10% PEG and 20 μM ThT. Results are mean ± SEM. One-way ANOVA; ****P ≤ 0.0001.

We then aimed to investigate the ability of ThT to detect fibrillar species formed within α-synuclein condensates (**Figure 1** and **Supplementary Movie 3**). To this end, monomeric α-synuclein labelled with the fluorophore Alexa Fluor 647 was incubated in the presence of 20 μM ThT, and observed for about 20 min (**Figure 1A**). This timeframe was chosen based on the observation that condensates transition from a liquid to a solid state about 20 min after the onset of liquid-liquid phase separation (**Supplementary Figures 1** and **2**). We observed a general increase in ThT fluorescence over time at various concentrations of α-synuclein, suggesting that fibrillar species had formed within the condensate (**Figure 1B,C**). The quantification of ThT emission resulted in a typical sigmoidal growth curve, suggesting that the formation of amyloid fibrils within the condensates follows a nucleationbased growth model of aggregation^10,11^(**Figure 1C,D**). During the first few min from the start of the liquid-liquid phase separation process, the vast majority of α-synuclein was present in its monomeric state (**Figure 1B,C**). After 10 min, a notable increase in ThT emission was detected (**Figure 1B,C**), as evidenced by an exponential growth phase, which is indicative of a rapid increase in cross-β α-synuclein species within condensates^37^. Such a speedy amplification of aggregate mass points towards the presence of secondary pathways, driving the amplification of cross-β aggregates^10,11,37^. Eventually, after approximately 18 min, the ThT emission reached a plateau, as most monomeric α-synuclein had been converted into fibrils (**Figure 1B,C**), indicating completion of the aggregation process.

While the presence of ThT-positive species indicates the presence of amyloid aggregates, it is also important to characterise their morphology at the endpoint of the assay. To this end, we performed transmission electron microscopy (TEM) imaging 24 min post the onset of liquid-liquid phase separation by obtaining a ThT-verified sample drop from a microscopy dish. As expected, the TEM images indicated that α-synuclein fibrils had indeed formed, which exhibited a cross-β-like network structure. Importantly, no major fibrillar structures were observed prior to liquid-liquid phase separation, and when individual components of the assay were assessed individually (**Supplementary Figure 4A,B**), which further confirms that our assay is capable of following the formation of α-synuclein fibrils following liquid-liquid phase separation.

### The ThT-based aggregation assay is sensitive to changes in liquid-liquid phase separation behaviour

Liquid-liquid phase separation is known to be driven by low-affinity interactions such as electrostatic and hydrophobic effects^38,39^. We used these observations to test the sensitivity of the aggregation assay described above in detecting changes in the protein interactions governing liquid-liquid phase separation. A ionic compound, sodium chloride (NaCl), and an aliphatic alcohol, 1,6-hexanediol, were chosen as controls due to their wide applications as effective tools to probe the material properties protein condensates and inclusions^19,40^. NaCl alters the electrostatic interactions between protein molecules in the liquid-liquid phase separation process, while 1,6-hexanediol works by interfering with the weak hydrophobic interactions that promote the formation of protein condensates.

As expected, 1,6-hexanediol had a concentration-dependent inhibitory effect on the formation of α-synuclein condensates (**Figure 2A,B**). While this finding shows that 1,6-hexanediol effectively reduced the maturation of the protein condensates and the formation of amyloid aggregates, it also emphasises that the condensate dissolution through this type of mechanism may not be a practical therapeutic strategy, as α-synuclein condensate formation was disrupted, and amorphous structures were predominantly formed at high 1,6-hexanediol concentrations (**Figure 2A**). More importantly, the fact that this compound may prevent α-synuclein from accessing a potentially physiological condensate state is likely undesirable. On the other hand, an increase in ionic strength led to an acceleration in α-synuclein aggregation (**Figure 2A,C**), a result that implicates the involvement of the acidic C-terminal region of α-synuclein in its phase behaviour. Taken together, these results demonstrate a reliable method of assessing α-synuclein aggregation within dense liquid condensates.

**Figure 2.**
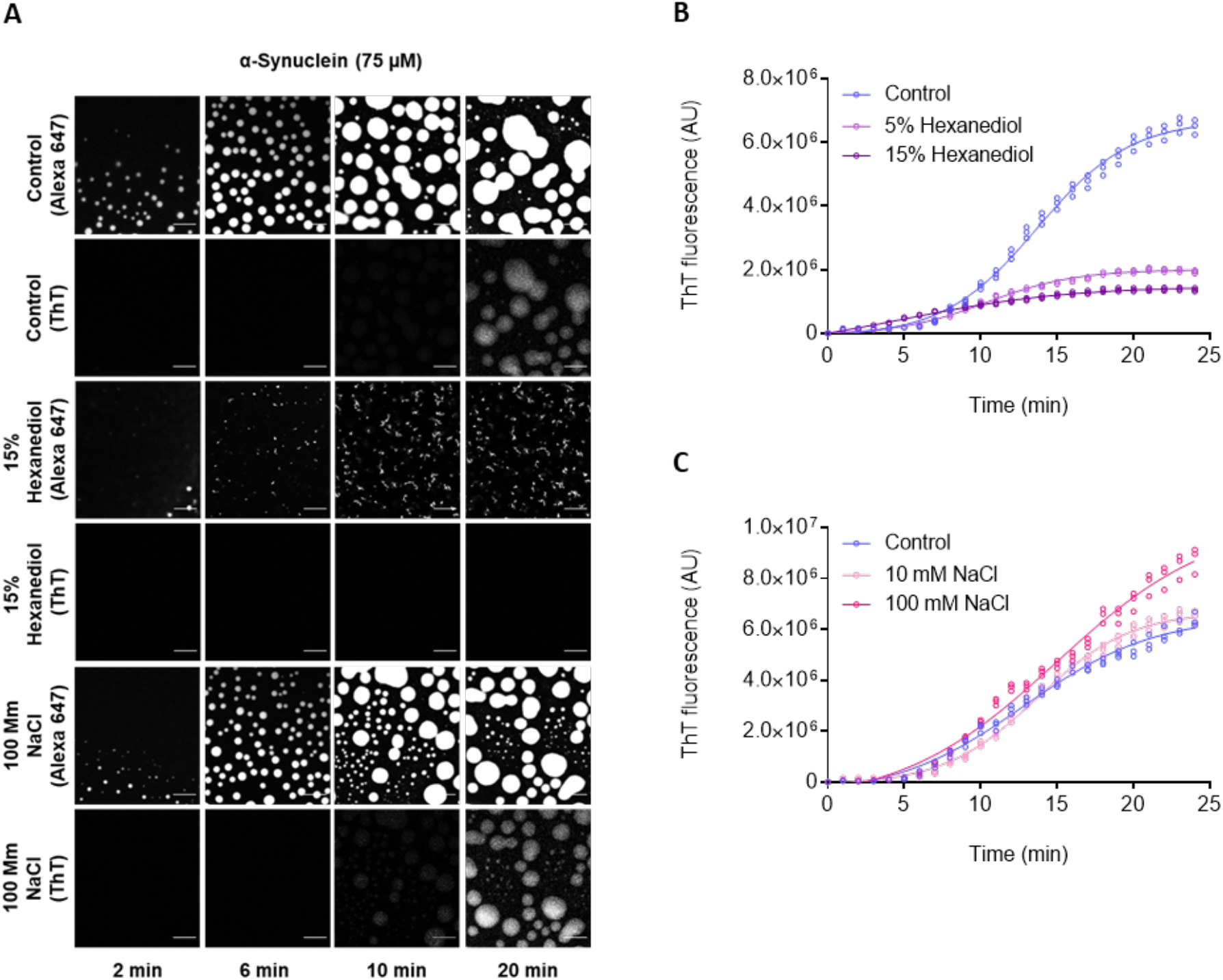
The ThT-based aggregation assay is sensitive to changes in liquid-liquid phase separation behaviour. **(A)** Images showing α-synuclein condensate formation (using Alexa Fluor 647 fluorescence) and aggregation (using ThT fluorescence) in the presence and absence (control) of 1,6-hexanediol (15% w/v), and of 100 mM NaCl. The images suggest that the assay is sensitive to changes in the interactions between α-synuclein monomers as a result of the presence of 1,6-hexandiol and sodium chloride. Scale bars represent 10 μm. **(B)** Quantification of the ThT emission over time for the images shown in panel A for 75 μM α-synuclein (control) (blue) in the presence of 15% (w/v) (dark purple) and 5% (w/v) (light purple) 1,6-hexanediol over a 24-min time period. **(C)** Rate of α-synuclein aggregation for the images shown in panel A in the presence and absence (blue) of 10 mM (light orange) and 100 mM (orange) NaCl over a 24-min time period. The curves represent a non-linear fit of the ThT signal (integrated density) over time. All experiments were performed using 75 μM α-synuclein in 50 mM Tris-HCl at pH 7.4 in the presence of 10% PEG and 20 μM ThT.

### A framework to elucidate the mechanism of α-synuclein aggregation within condensates

Using the assay described above, we aimed to unravel the kinetic mechanisms by which monomeric α-synuclein transitions into fibrillar aggregates within liquid condensates. We first set out to establish a model of α-synuclein aggregation within condensates in an unperturbed system. Using a web-based software, AmyloFit we can characterise the complex reaction networks associated with protein aggregation using experimentally-obtained kinetic data^37^. AmyloFit normalises the aggregation data and plots them as a function of initial monomer concentration (m_0_) to extract rate constants and scaling exponents, which reveal the dominant aggregation mechanisms.

When trying to obtain model of α-synuclein aggregation within condensates in the unperturbed system, we were unable to attain a global fitting and scaling exponents for our data. We rationalised these findings based on the dynamic nature of the exchange of monomeric proteins between the dilute and dense phase, which implies that variations in protein concentrations in the dilute phase yield similar protein concentrations within the dense phase (**Figure 3**). Analysis of individual sigmoidal curves, however, does not contain enough information for the different underlying molecular mechanisms to be distinguished as they can be fitted by a range of different reaction schemes (**Figure 3B**). Nevertheless, the sigmoidal ThT curves that we obtained show a lag phase followed a growth phase and a plateau phase, therefore presence of fibril-dependent secondary processes can be assumed.

**Figure 3.**
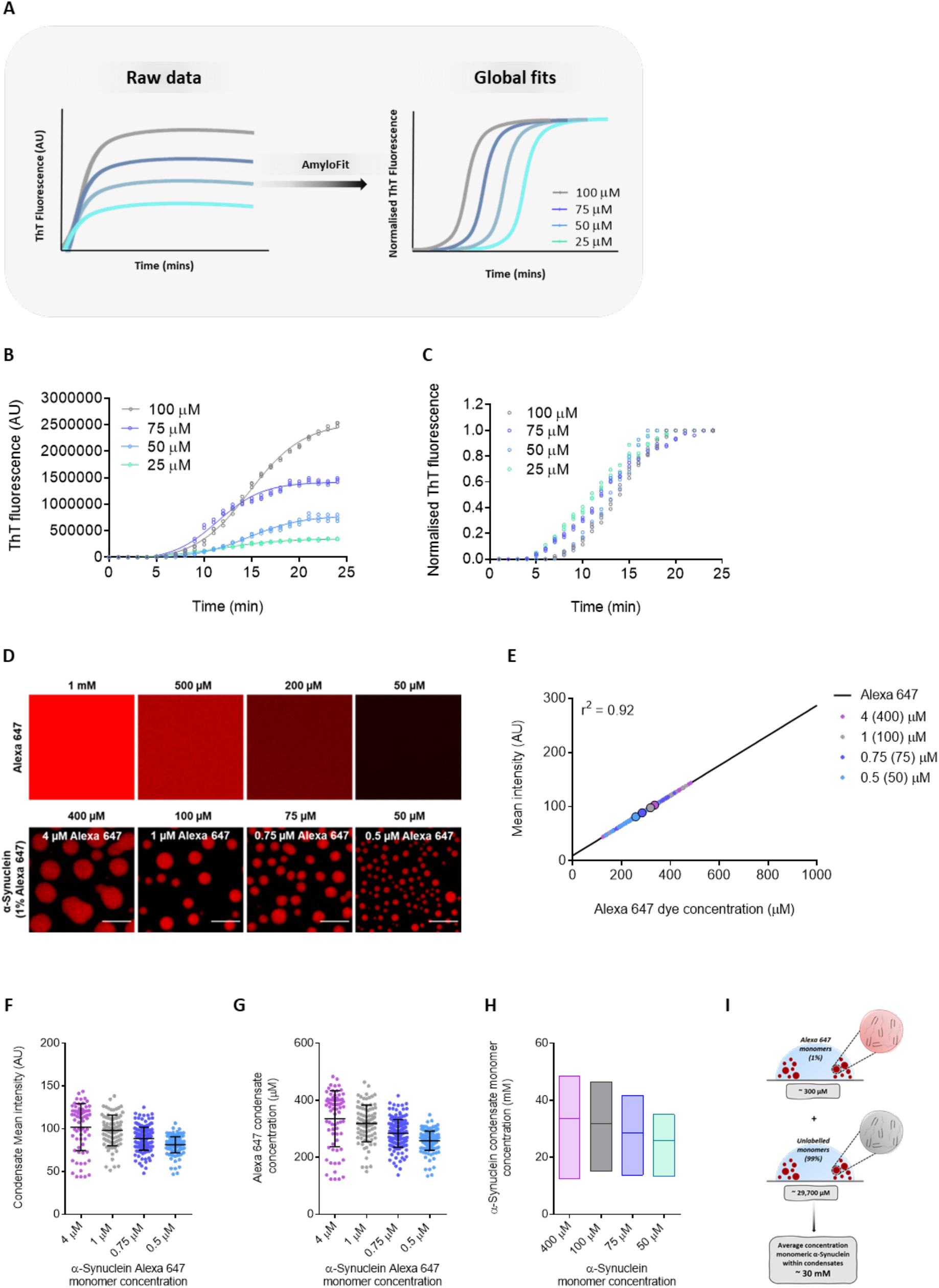
The kinetics of α-synuclein α-synuclein within liquid condensates show very weak concentration dependence. **(A)** ThT emission curves obtained through an aggregation assay. In this type of aggregation assay, ThT fluorescence readings at different protein concentrations are processed and normalised to obtain a global fitting of a kinetic model. **(B-C)** Within condensates, normalised aggregation data at four α-synuclein concentrations (100 μM (grey), 75 μM (blue), 50 μM (cyan) and 25 μM (turquoise)), yielded almost no variations between different concentrations. **(B)** Quantification of ThT emission over a 24-min time period for the four different α-synuclein concentrations. **(C)** Normalised representative aggregation kinetic traces of α-synuclein ThT emission shown in panel B. All experiments were performed in 50 mM Tris-HCl at pH 7.4 in the presence of 10% PEG and 20 μM ThT. **(D)** Fluorescence images of Alexa Fluor 647 at a range of concentrations (1000, 500, 200 and 50 μM) and images of monomeric α-synuclein (400, 100, 75 and 50 μM) condensates 10 min from the onset of liquid-liquid phase separation. Scale bar represents 10 μm. **(E)** Fluorescence intensity of Alexa Fluor 647 from at a range of 1-1000 μM from images shown in panel B were used to obtain a standard curve of fluorescence signal for calibration (linear regression, r^2^ = 0.92). The small circles are the individual condensates measured for each α-synuclein concentration (n > 70 condensates per concentration) and the big bold circles indicate the mean α-synuclein condensate intensity for each concentration (400 μM (light purple), 100 μM (grey), 75 μM (blue) and 50 μM (cyan) and 4 μM, 1 μM, 0.75 μM and 0.5 μM for their respective 1% Alexa Fluor 647 protein concentration). **(F)** Quantification of the mean fluorescence intensity for individual condensates at each α-synuclein Alexa Fluor 647 concentrations 10 min from the onset of liquid-liquid phase separation. **(G)** The concentration within liquid droplets of α-synuclein labelled with Alexa Fluor 647 at a range of bulk concentrations (400, 100, 75 and 50 μM) was estimated to be 335±98, 319±64. 284±48 and 258±34 μM after 10 min from the onset of liquid-liquid phase separation (n > 70 condensates per concentration). **(H)** The total α-synuclein concentration (labelled and unlabelled) within condensates was estimated at an average of 33.5, 31.9, 28.4 and 25.8 mM for samples at 400, 100, 75 and 50 μM α-synuclein concentration (labelled and unlabelled), respectively. **(I)** Schematic illustration of the estimated total α-synuclein concentration within condensates, which is approximately 30 mM from the results in panel H. Data are shown for a representative experiment that was repeated three times with similar results. All liquidliquid phase separation experiments were performed in 50 mM Tris (pH 7.4) and 10% PEG.

Given that the study of the mechanisms of α-synuclein aggregation using AmyloFit generally involves the use of experimental data obtained from ThT assays conducted in bulk solution (**Figure 3A**), we aimed to see if we could obtain the model of α-synuclein aggregation at the condensates through a different method. To do this, we determined the monomeric protein concentration within the condensates as a function of time and of the concentration in the dilute phase. The fluorescence intensity of Alexa Fluor 647 was used as readout for monomeric protein concentrations, as this accounts for 1 molar % of the total α-synuclein concentration (**Figure 3**). We observed that the monomeric protein concentration within condensates, 6 min post liquid-liquid phase separation did not change despite an increase in amyloid formation (**Supplementary Figure 6A**). As condensates mature and grow in a concentration-dependent manner, we wanted to establish if there was a correlation between the condensate size and the monomeric concentration of α-synuclein in the dense phase. At all tested concentrations, there was a weak correlation between these variables, and the differences between concentrations were not significant (**Supplementary Figures 5** and **6B**). Given these results, we then estimated the total concentration of α-synuclein (Alexa Fluor 647 labelled (1%) and unlabelled (99%)) in liquid droplets to be about 30 mM for all concentrations, by using a previously described method where the fluorescence intensity of individual α-synuclein condensates was measured and Alexa Fluor 647 was used to make a standard curve for fluorescence intensity^34^ (**Figure 3**). Thus, the estimated total concentration of α-synuclein in condensates is about three orders of magnitude higher than in the diluted phase.

### Secondary processes dominate α-synuclein aggregation within condensates

We next investigated the role of aggregate-dependent secondary processes on α-synuclein aggregation within condensates. The addition of preformed fibrils at the beginning of the aggregation assay is typically used as a qualitative way to check for the presence of secondary processes, as they act as seeds to aid in aggregate growth and multiplication^37^. On these grounds, we aimed to bypass primary nucleation (with rate constant *k_n_*) events by adding a small amount of seeds (2%), where the rate of fibril formation is driven by secondary nucleation (with rate constant *k_2_*), fragmentation (with rate constant *k*_-_) and elongation (with rate constant *k*_+_)^15^. The addition of large amounts (25%) of preformed seeds leads to primary and secondary nucleation being bypasses and elongation of the preformed fibrils being the dominant process of aggregation^15^ (**Figure 4A**). If secondary nucleation is indeed present in α-synuclein aggregation within condensates, we expect a shortening in the lag phase in the presence of the preformed fibrils (**Figure 4A**). Our results indicate that α-synuclein aggregation within condensates was accelerated in the presence of seeds as most newly formed fibrils were generated through secondary nucleation (**Figure 4D**). In the presence of 25% preformed fibrils, α-synuclein aggregation was accelerated drastically to the extent that the plateau phase was reached 10 min post liquid-liquid phase separation (**Figure 4D,F**). Visually, the presence of preformed fibrils within our assay was observed in the dilute phase prior to liquid-liquid phase separation (**Figure 4B**). Upon liquid-liquid phase separation, preformed fibrils were observed to colocalise with liquid droplets, thus speeding up amyloid formation (**Figure 4B,C**). Overall, the data suggest that secondary processes are involved in the aggregation of α-synuclein within condensates. However, to confirm this result, the concentration of preformed fibrils within the liquid droplets needs to be calculated.

**Figure 4.**
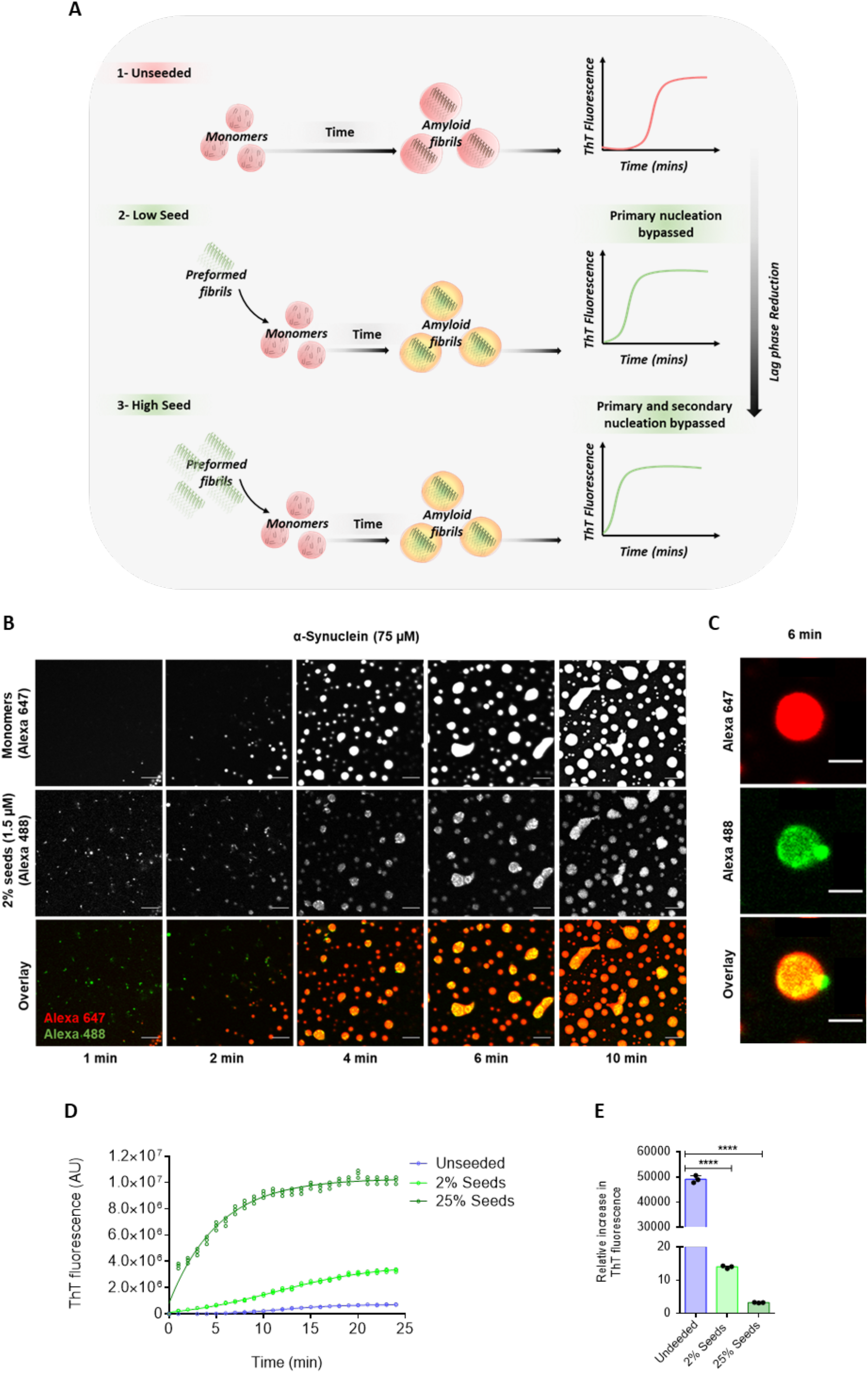
α-Synuclein primary nucleation within condensates is bypassed by preformed fibrils. **(A)** Schematic diagram illustrating the seeding process within the condensates. In the presence preformed fibrils (seeds), aggregation is accelerated (2 and 3), whereas in the absence of seeds, formation of amyloid-containing condensates is slower as a function of time (1). Increasing seed concentration results in reduced lag phase. **(B)** Representative fluorescence imaging displaying α-synuclein monomer (75 μM) condensate formation (Alexa Fluor 647) in the presence of 2% (1.5 μM) Alexa Fluor 488-labelled preformed fibrils, 10% PEG in 50 mM Tris-HCl (pH 7.4). At 1 min the presence of preformed fibrils within the droplet are observed in the dilute liquid phase. Upon the formation of liquid droplets, preformed fibrils are seen to co-localise with condensates thereby accelerating aggregation by propelling the formation of amyloid-containing condensates. Scale bar represents 10 μm. **(C)** Image of a liquid droplet showing in which preformed fibrils (Alexa Fluor 488) are directly recruited into α-synuclein condensate (Alexa Fluor 647). Scale bar represents 10 μm. **(D)** Progression of ThT emission as a function of time (24 min) for 75 μM α-synuclein monomers (unseeded) (blue) with the addition of 2% (1.5 μM) (light green) and 25% (18.75 μM) (dark green) preformed fibrils. The curve represents a non-linear fit of the ThT signal (integrated density) over time. **(E)** Relative increase in the raw data values of the ThT fluorescence intensities from time from the onset of liquid-liquid phase separation to 24 min post it. All aggregation assays were performed in 50 mM Tris-HCl (pH 7.4) in the presence of 10% PEG and 20 μM ThT. Results are mean ± SEM. One-way ANOVA; ****P ≤ 0.0001.

A standard liquid-liquid phase separation assay without ThT was employed to determine the concentration of preformed fibrils within the condensates. This assay was conducted using Alexa Fluor 488 labelled preformed fibrils, which were observed to colocalise with the liquid droplets 6 min post liquid-liquid phase separation (**Figure 4B,C**). Bearing this in mind, several fluorescently labelled preformed fibrils containing condensates were assessed for their fibrillary content from several time points between 6-20 min. As a result, we estimated the concentration of preformed α-synuclein fibrils within liquid droplets to be 37 μM (range: 27-75 μM) for 2% seeds (1.5 μM), where the intensity of individual α-synuclein condensates (75 μM) containing preformed fibrils (25-100 μM) was measured, and a range of Alexa Fluor 488 fibril standards (50-300 μM) was used to create a standard curve of Alexa Fluor 488 mean fluorescence intensity (**Figure 5**). With regards to 25% seeds (18.75 μM) the average mean fluorescence intensity was greater than that of the highest concentration (300 μM) used for the standard curve. In view of this, the equation *c^I^* ≅ 24.2*c^tot^* was derived to estimate the fibril concentration inside the condensate. Here, 24.2 is the constant obtained from the slope of the standard curve, *c^I^* refers to the fibrils partitioned inside the condensate, where as *c^tot^* refers to the total fibril concentration added to the system (1.5 and 18.75 μM) (**Figure 5C**). Using this equation, the concentration of preformed fibrils within condensates at 25% seeds was estimated to be at 454 μM, which was further validated through experimental analysis (**Figure 5B,D**).

**Figure 5.**
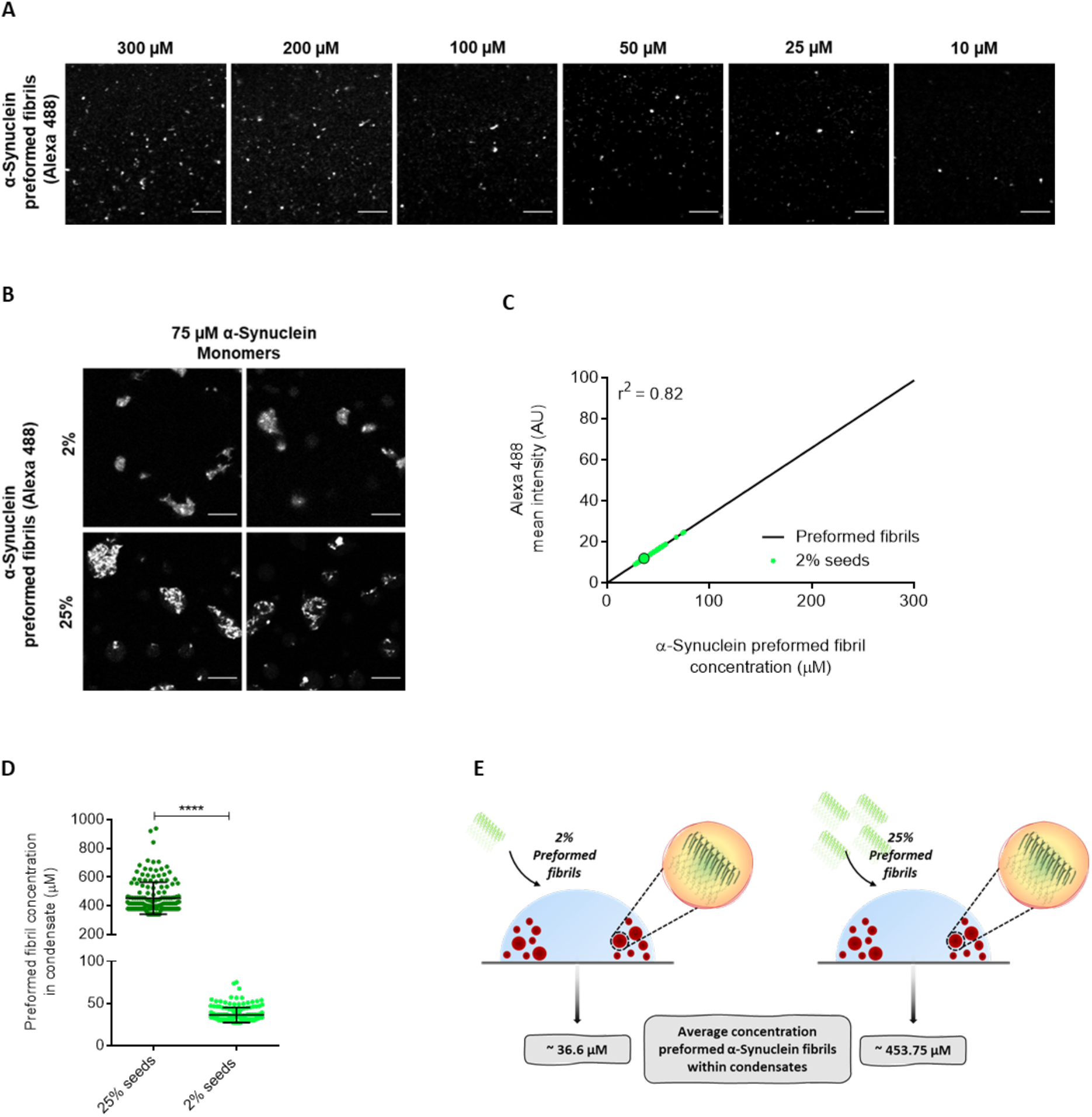
Determination of α-synuclein preformed fibril concentration within condensates. **(A)** Fluorescence images of Alexa Fluor 488 labelled preformed fibrils at a range of concentrations (300, 200, 100, 50, 25 and 10 μM). Scale bar represents 10 μm. **(B)** Representative fluorescence imaging displaying co-localisation of 2% and 25% preformed fibrils (Alexa Fluor 488) within α-synuclein condensates (Alexa Fluor 647) 10 min post liquid-liquid phase separation). Scale bar represents 10 μm. **(C)** Alexa Fluor 488 labelled preformed fibrils at a range of 10-300 μM from images shown in (A) were used for a standard curve of Alexa Fluor 488 fluorescence signal (linear regression, r^2^ = 0.82). The small circles are the individual condensates (n > 100 condensates) measured 10 min from the onset of liquid-liquid phase separation and the big bold circle indicates the mean of all α-synuclein fluorescently labelled preformed fibril containing condensate intensity. Fluorescence signal of condensates with assay containing 25% preformed fibrils had a higher 488 intensity than the intensity of 300 μM preformedfibrils usedfor standard curve. **(D)** Concentration of 2% (1.5 μM) and 25% (18.75 μM) Alexa Fluor 488 labelled α-synuclein preformed fibrils within liquid droplets of α-synuclein (75 μM) was estimated at an average of 36 μM (SD = ±9, Min = 27 μM, Max = 75 μM) and 454 μM (SD = ±112, Min = 341 μM, Max = 939 μM) respectively. 10 min from the onset of liquid-liquid phase separation (n > 100 condensates). **(E)** Summary schematic illustrating the total concentration of α-synucleinpreformed fibrils within condensates to be approximately 36.6 μM and 453.75 36.6 μM for 2% and 25 % seeds, respectively. Data are from a representative experiment that was repeated three times with similar results. All seeded aggregation experiments were performed in 50 mM Tris (pH 7.4) and 10% PEG. Results are mean ± SEM. One-way ANOVA; ****P ≤ 0.0001.

Having obtained the concentrations of preformed fibril within condensates, we tested the fit of the seeded kinetics data to three different models (**Figure 6**). The first fit clearly shows that the aggregation data are not consistent with a model that includes only primary nucleation and elongation. In contrast, the models with secondary processes resulted in good fits (**Figure 6**). As primary nucleation and secondary nucleation have a different concentration dependence, if we assume that fragmentation does not contribute significantly to aggregate proliferation under the conditions used, the critical fibril concentration at which secondary nucleation becomes the dominant mechanism is 20 mM in our system (ratio of the primary to secondary nucleation rate constants, *K_n_/K_2_m_o_*; the parameters obtained in **Figure 6**).

**Figure 6.**
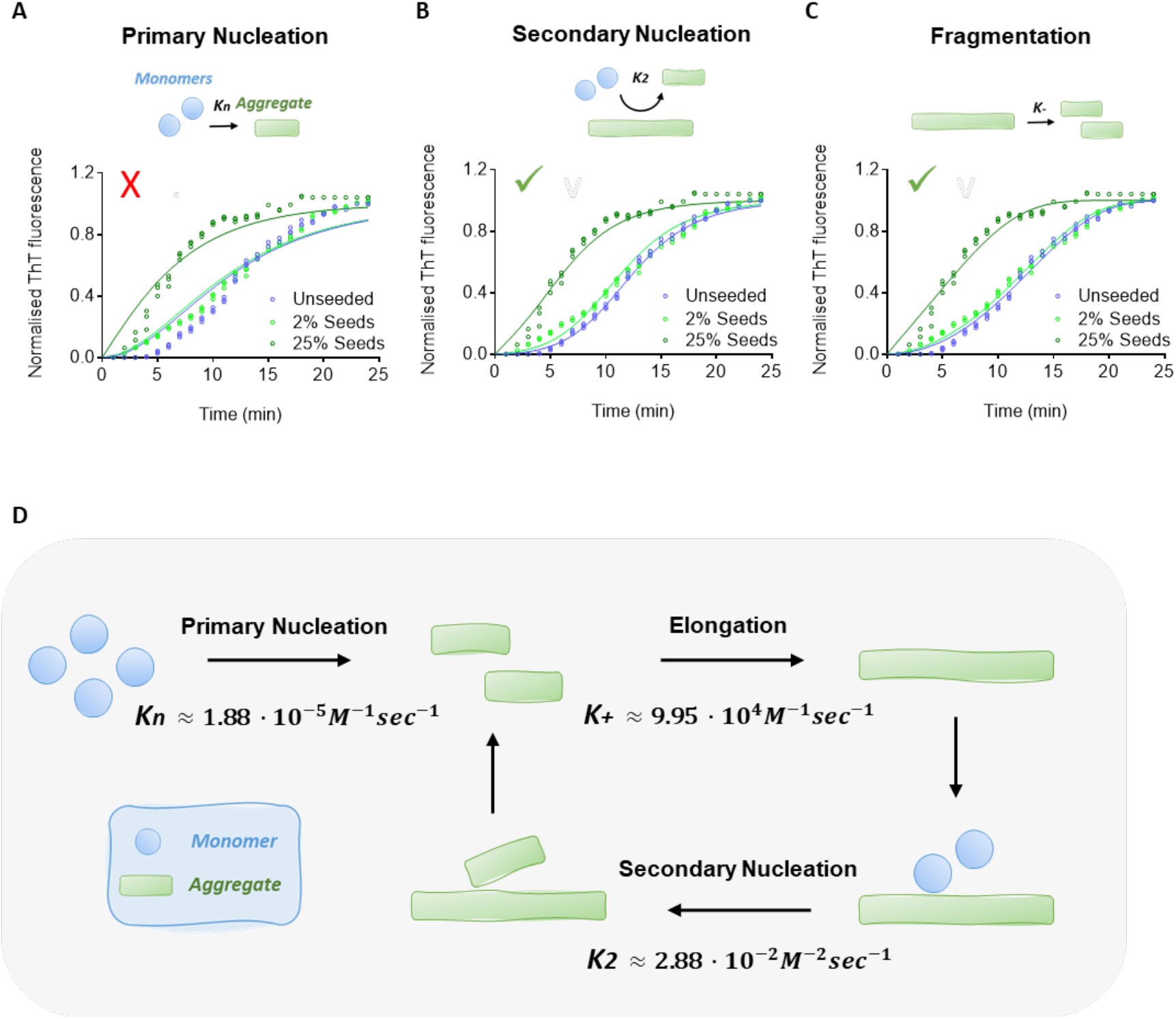
Secondary processes dominate α-synuclein aggregate proliferation within liquid condensates at physiological pH. The data were analysed using AmylioFit, with the normalised aggregation curves measured as a function of time in the ThT aggregation assay from the unperturbed system (blue), as well as in the presence of 2% (light green) and 25% (dark green) seeds. The graphs represent the best fits of a model where primary nucleation **(A)**, secondary nucleation **(B)** and fragmentation **(C)** is described to be the dominant mechanism of aggregation. The mean residual errors (MREs) were 0.009, n_c_ = 2 **(A)**, 0.002, n_c_ = 2, n_2_ = 1 **(B)**, 0.002, n_c_ = 2 **(C)**. Solid lines represent the best fit to each respective condition. An m_0_ = 300 mM was used for all fittings, for the unperturbed system M_0_ = 0. For seeded assay M_0_ = 36.3 μM and 453.75 μM for 2% and 25% seeds, respectively. All experiments were performed three times using 75 μM α-synuclein in 50 mM Tris-HCl (pH 7.4) in the presence of 10% PEG and 20 μM ThT. Using the assay reported in this work, we illustrate the microscopic processes involved in α-synuclein aggregation and the associated rate constants, in the case when fragmentation is not contributing significantly. Despite the low values of the rate constants for primary nucleation and secondary nucleation, the aggregation process proceeds rapidly within condensates because of the high concentration of the protein in this phase.

## Conclusions

We have reported a fluorescence-based aggregation assay to determine the microscopic processes governing the aggregation of α-synuclein within liquid condensates (**Figure 6**). Our findings have revealed that α-synuclein can undergo spontaneous primary nucleation and rapid aggregate-dependent aggregate proliferation at physiological pH. We anticipate that this approach will enable further studies to establish whether α-synuclein aggregation via the liquid-liquid phase separation pathway is relevant to the onset and progression of Parkinson’s disease. In perspective, the possibility of studying the aggregation process of α-synuclein under physiological conditions with the fast and reliable assay that we have reported will open novel opportunities for screening compounds to inhibit α-synuclein aggregation and its pathological consequences in Parkinson’s disease and related synucleinopathies^41,42^.

## Materials and Methods

### Purification and labelling of α-synuclein

Human wild-type α-synuclein and its cysteine variant A90C were expressed and purified as previously described, using *E. coli* BL21 (DE3)-gold competent cells (Agilent Technologies) transformed with the pT7-7 plasmid encoding α-synuclein; expression was induced by the addition of 1 mM isopropyl β-d-1-thiogalactopyranoside (IPTG)^43^. Following purification, proteins were concentrated, and buffer exchanged into 50 mM trisaminomethane-hydrochloride (Tris-HCl) at pH 7.4 using Amicon Ultra-15 Centrifugal Filter Units (Merck Millipore). The A90C variant was labelled with 1.5-fold molar excess of C5 maleimide-linked Alexa Fluor 647 (Invitrogen Life Technologies) overnight at 4 °C under constant gentle stirring. The unbound dye was removed using Amicon Ultra-15 Centrifugal Filter Units and buffer exchanged into 50 mM Tris-HCl at pH 7.4. The final protein concentration was measured using ultraviolet-visible (UV-vis) spectroscopy on a Cary 100 system (Agilent Technologies). All proteins were aliquoted, flash-frozen in liquid nitrogen, stored at −80 °C and thawed once before each experiment.

### Preparation and labelling of preformed fibrils of α-synuclein

α-Synuclein seed fibrils were formed from recombinant α-synuclein wild-type or cysteine variant (A90C) monomers diluted in buffer (50 mM Tris-HCl at pH 7.4) to concentrations of approximately 500 μM. Monomers in microcentrifuge Eppendorf tubes (Axygen low-bind tubes) were incubated at 40 °C, with constant stirring speed (1,500 rpm) with a teflon bar and left to fibrillate on an RCT Basic Heat Plate (RCT Basic, model no. 0003810002; IKA, Staufen, Germany) for up to 72 h. Samples were centrifuged (Centrifuge 5427 R) at 4 °C, 14,000 rpm for 15 min. The supernatant was used to determine the monomer concentration and fibril concentration was determined as monomer equivalents. For visualisation purposes, A90C α-synuclein fibrils were labelled with 1.5-fold molar excess of C5 maleimide-linked Alexa Fluor 488 (Invitrogen Life Technologies) overnight at 4 °C under constant but gentle stirring. The excess dye was removed by at least 3 sequential 50 mM Tris-HCl washes by centrifugation (4 °C, pH 7.4, 14,000 rpm) for 15 min. Before each experiment, seed fibrils were pretreated by 15 sec (15 pulses) sonication at 10% power with 50% duty cycle using a Microtip sonicator (Bandelin Sonopuls HD2070) to disperse lumped fibrils.

### α-Synuclein liquid-liquid phase separation assay

To induce liquid droplet formation, non-labelled wild-type α-synuclein was mixed with the A90C variant labelled with Alexa Fluor 647 at a 100:1 molar ratio in 50 mM Tris-HCl, (pH 7.4) and 10% polyethylene glycol 10,000 (PEG) (Thermo Fisher Scientific) at room temperature (20-22 °C). Additionally, NaCl, 1,6-hexanediol and dimethyl sulfoxide (DMSO) and preformed fibrils were supplemented in the liquid-liquid phase separation assay. 10 μL of the final mixture was pipetted on a 35 mm glass bottom dish (P35G-1.5-20-C, MatTek Life Sciences) and immediately imaged on a Leica TCS SP8 inverted confocal microscope using a 60×/1.4 HC PL Apo CS oil objective (Leica Microsystems). The excitation wavelength was set to 633 nm for all experiments. All images were processed and analysed in ImageJ (NIH).

### α-Synuclein aggregation assay within dense liquid condensates

20 μM ThT (Sigma) and additional compounds and preformed fibrils, depending on the experiments, were mixed with monomeric wild-type α-synuclein, containing 1 molar % A90C α-synuclein labelled with Alexa Fluor 647 prior to each experiment. The assay mixture, which included 50 mM Tris-HCl, pH 7.4, 10% PEG 10,000, was pipetted on to a 35 mm glass bottom dish and imaged on a Leica TCS SP8 inverted confocal microscope using a 60×/1.4 HC PL Apo CS oil objective. The excitation wavelength 633 nm and 488 nm were used for Alexa Fluor 647 labelled α-synuclein and ThT, respectively. Images were processed in ImageJ; an area adjacent to the edge of the droplet was cropped and analysed, thus allowing for time-dependent measurement of amyloid formation at the droplets.

### Transmission electron microscopy (TEM)

α-Synuclein samples from the ThT-based aggregation assay were obtained prior to fibril formation mediated by liquid-liquid phase separation and after fibril formation. The obtained sample (5 μL) was deposited on a carbon film of 400 mesh 3 mm copper grid. The grids were washed once with Milli-Q water, then incubated with 1% (w/v) uranyl acetate for 2 min and washed twice again with Milli-Q water before being air-dried at room temperature. Samples were imaged at the Cambridge Advanced Imaging Centre using a Tecnai G2 transmission electron microscope operating at 80-200 keV.

### Estimation of α-synuclein concentration in liquid droplets

The fluorescence standard was performed by imaging solutions of the Alexa Fluor 647 dye at 1000, 500, 200, 100, 50, 20 and 1 mM in 50 mM Tris-HCl at pH 7.4 with the addition of 10% PEG. Using the same acquisition settings, images were taken of α-synuclein condensates at 400, 100, 75, 50 and 25 μM 10 min after liquid-liquid phase separation to estimate the local concentration of α-synuclein liquid droplets. Several condensates (approximately 70 per concentration) were analysed by drawing a circular region of interest around the condensate to obtain the average pixel intensity for each droplet. Linear regression analysis of the intensity of Alexa Fluor 647 standard images (r^2^ = 0.92) was used to interpolate the estimated concentration of α-synuclein proteins within condensates.

### Estimation of α-synuclein preformed fibril concentration in liquid droplets

An Alexa Fluor 488 standard was carried out using various concentrations of preformed fibrils of A90C α-synuclein labelled with Alexa Fluor 488 in 50 mM Tris-HCl at pH 7.4 with the addition of 10% PEG. Using the same acquisition settings, images were taken of α-synuclein condensates 6 min after liquid-liquid phase separation at 75 μM containing 2% and 25% Alexa Fluor 488 labelled preformed fibrils to estimate the local concentration of preformed fibrils within the liquid droplet. Several condensates (approximately 100) were analysed by drawing a circular region of interest around the condensate to obtain the average pixel intensity for each droplet. Linear regression analysis of the intensity of Alexa Fluor 488 fibrils standard images (r^2^ = 0.82) was used to estimate the concentration of α-synuclein preformed fibrils within condensates.

### Equation for the estimation of α-synuclein fibril concentration inside droplets

If we consider a system of total volume *V* and containing a dense phase (droplet phase) coexisting with a dilute phase, the dense phase (which we term phase I) occupies a volume *V^I^*, while the volume of the dilute phase (which we term phase II) is *V^II^* = *V* — *V^I^*, such that

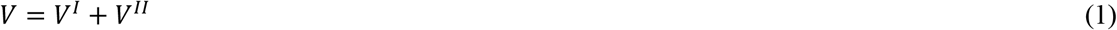

If we add to this system a total concentration *c^tot^* of fibrils, these fibrils will partition differently in phases I and II depending on their relative interaction with the dense and dilute phases. We capture this behaviour by means of the partitioning degree Γ, defined as

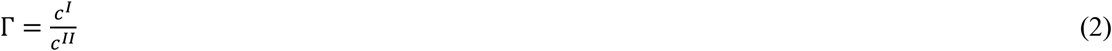

While Γ can be determined from the Flory-Huggins of phase separation^44^, we will consider it as a known parameter in our calculations, where *c^I^* and *c^II^* are the concentrations of fibrils in phases I and II, respectively. The total fibril concentration in the system is the average of the concentrations *c^I^* and *c^II^* weighted by the volumes of the respective phases

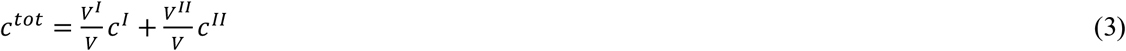

This condition states that the total number of fibrils in the system is conserved. By combining Eqs. (1), (2) and (3) we can solve for *c^I^* and *c^II^* and find

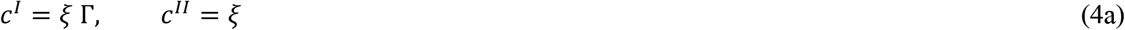

where

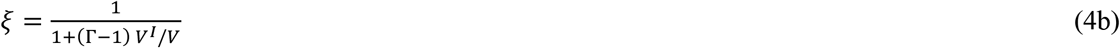

Using Eq. (4) we can calculate the concentration of fibrils in the droplet phase from *c^tot^*, Γ and *V^I^/V*. In the limit when Γ → ∞ (when the fibrils partition mostly inside the droplet phase), Eq. (4) reduces to

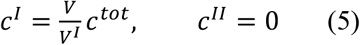

In this regime, we have a linear relationship linking *c^I^* to *c^tot^*. The slope of this relationship is *V/V^I^* and relates to the volume fraction *V^I^/V* occupied by the dense phase. In our experiments we measured *c^I^* = 36.6 μM for *c^tot^* = 1.5 μM and *c^I^* = 453.75 μM for *c^tot^* = 18.75 μM. With these data points we estimated a slope *V/V^I^* = 24.2.

### Image analysis of α-synuclein liquid droplets

Images of wild-type α-synuclein, containing 1 molar % α-synuclein Alexa Fluor 647 labelled A90C α-synuclein were acquired on a Leica TCS SP8 inverted confocal microscope system. All images within an experiment were acquired using identical confocal settings (scan speed, resolution, magnification, laser intensity, gain, and offset). Images were analysed by applying threshold functions in ImageJ software that identified the phase separated α-synuclein condensates and excluded the background of the image. All condensates within the threshold limits were analysed for total area (μm^2^), average size of individual droplets (μm^2^) and mean fluorescence intensity of individual droplets (arbitrary units) (**Figure S4**).

### AmyloFit data analysis

The aggregation of α-synuclein within droplets was monitored as described above. The experimental ThT fluorescence readout was uploaded on the free online platform AmyloFit^37^ (https://www.amylofit.ch.cam.ac.uk). Next, the software normalised the data to 0% and 100% by averaging the values at the baseline and the plateau of the reaction. Upon data normalisation, the concentration of aggregate mass could be observed as a function of time. An advanced basin-hopping algorithm was applied to fit the experimental data to a s model of protein aggregation. The initial monomer concentration (m_0_) was set to 30 mM. Reaction orders for primary and secondary nucleation were set to 2. The number of basin hops was set to 40 for each fit. The applied model was only considered suitable if it was able to match the experimental data well. The AmyloFit user manual can be consulted for more in-depth information on the fitting procedure^37^.

### Statistical analysis

All statistical analyses were performed in GraphPad Prism 8 (GraphPad Software). Data are presented as means ± SEM from at least 3 independent biological replicates, unless indicated otherwise. The statistical significance was analysed either by 2-tailed Student’s t test or one-way ANOVA followed by Bonferroni’s multiple-comparison.

### Online content

Any methods, additional references, Nature Research reporting summaries, source data, extended data, supplementary information, acknowledgements, peer review information; details of author contributions and competing interests; and statements of data and code availability are available at https://doi.org/xxx.

## Supplementary Information

### Supplementary Methods

#### Preformed fibril bound thioflavin T (ThT) assay

To study the concentration-dependent emission of fibril-bound ThT, wild-type α-synuclein preformed fibrils (100, 75, 50, 20 and 10 μM) were incubated with 20 μM ThT in 50 mM Tris-HCl and 10% PEG. For control monomeric wild-type α-synuclein were also treated with 20 μM ThT in 50 mM Tris-HCl and 10% PEG. 10 μL of the final mixture was pipetted on a 35 mm glass bottom dish and immediately imaged on a Leica TCS SP8 inverted confocal microscope using a 60×/1.4 HC PL Apo CS oil objective (Leica Microsystems). The excitation wavelength was 488 nM for all ThT-based experiments. All images were processed and analysed in ImageJ (NIH).

#### Fluorescence recovery after photobleaching (FRAP) experiment

FRAP experiments were performed on *in vitro* condensates using a Leica TCS SP5 inverted stage scanning confocal microscope. To conduct FRAP experiments, a 40x magnification oil objective (40x/1.3 HC PL Apo CS oil) was used. The argon laser was set to 30% power with bidirectional scanning speed of 400 Hz and the 488 nm laser line was set to 20% power in FRAP wizard. Time-lapse images of pre-bleached and bleached (ROI) condensate were collected at a 2 sec frame rate, recovery was monitored at a 20 sec frame rate. FRAP recovery data charts with intensity values averaged over the ROIs for all frames were evaluated and processed by the Leica TCS SP5 FRAP wizard system.

**Supplementary Figure 1.**
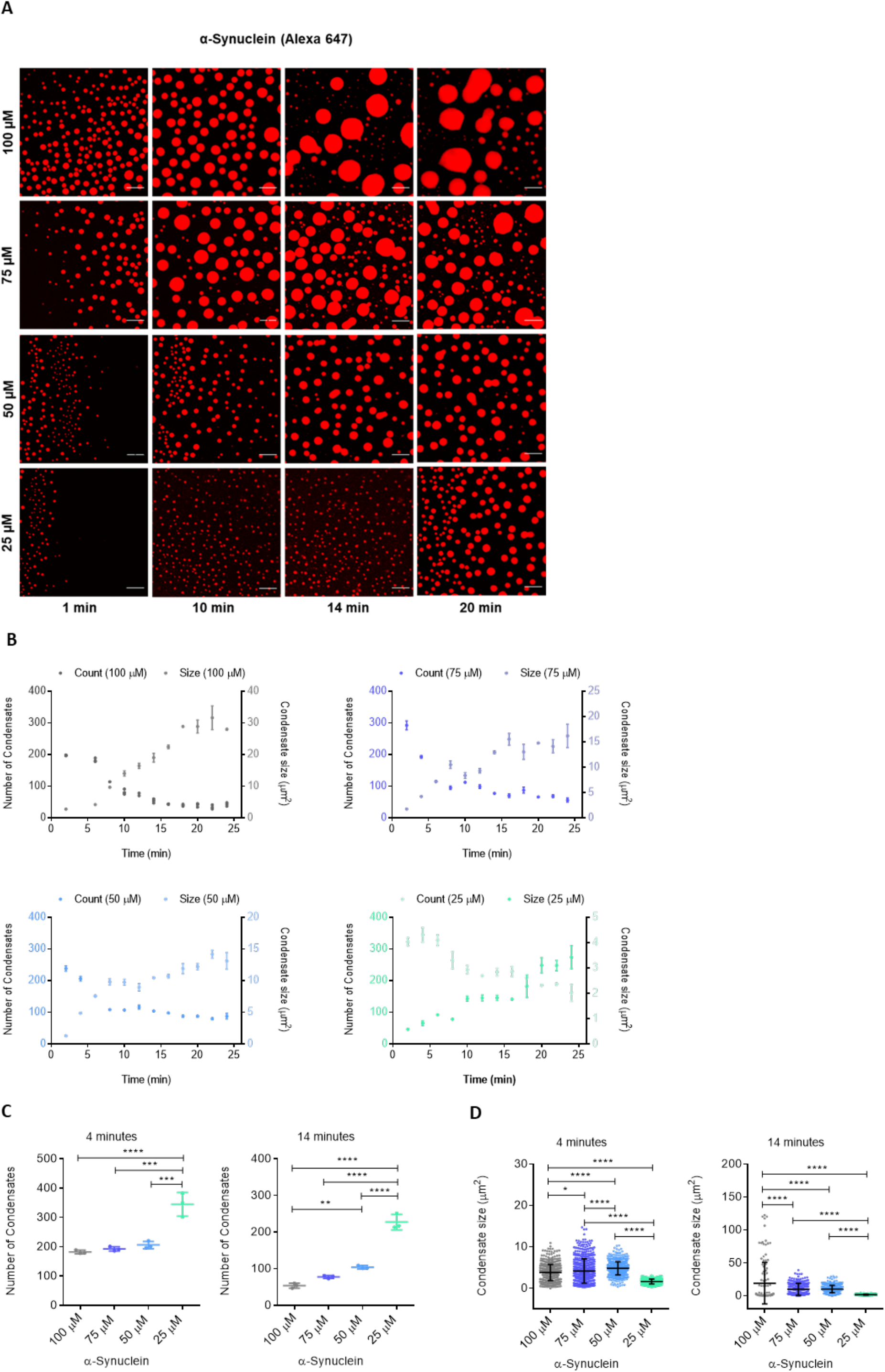
α-Synuclein forms liquid condensates at physiological concentrations under crowding conditions. **(A)** Fluorescence microscopy images of α-synuclein forming μm-sized condensates (labelled with Alexa Fluor 647) in the presence of crowding agent (10% PEG) in 50 mM Tris-HCl at physiological concentrations (100-25 μM) and pH (7.4) over time. The scale bar represents 10 μm. **(B)** Quantification of condensates number and size at indicated time points at physiological concentrations of 100 μM (grey), 75 μM (blue), 50 μM (cyan) and 25 μM (turquoise) of α-synuclein under crowded conditions in 50 mM Tris-HCl and pH 7.4. The inverse relation between number (dark colour) and size (light colour) illustrates the Ostwald ripening of the condensates over time. **(C,D)** Quantification of total number (C) and size distribution (D) of condensates at 4 and 14 min after liquidliquid phase separation. All experiments were performed in 50 mM Tris-HCl at pH 7.4 in the presence of 10% PEG. The data represent the mean ± SEM of n=3 individual experiments. One-way ANOVA. *P < 0.05, **P < 0.01, ***P < 0.001, ****P < 0.0001.

**Supplementary Figure 2.**
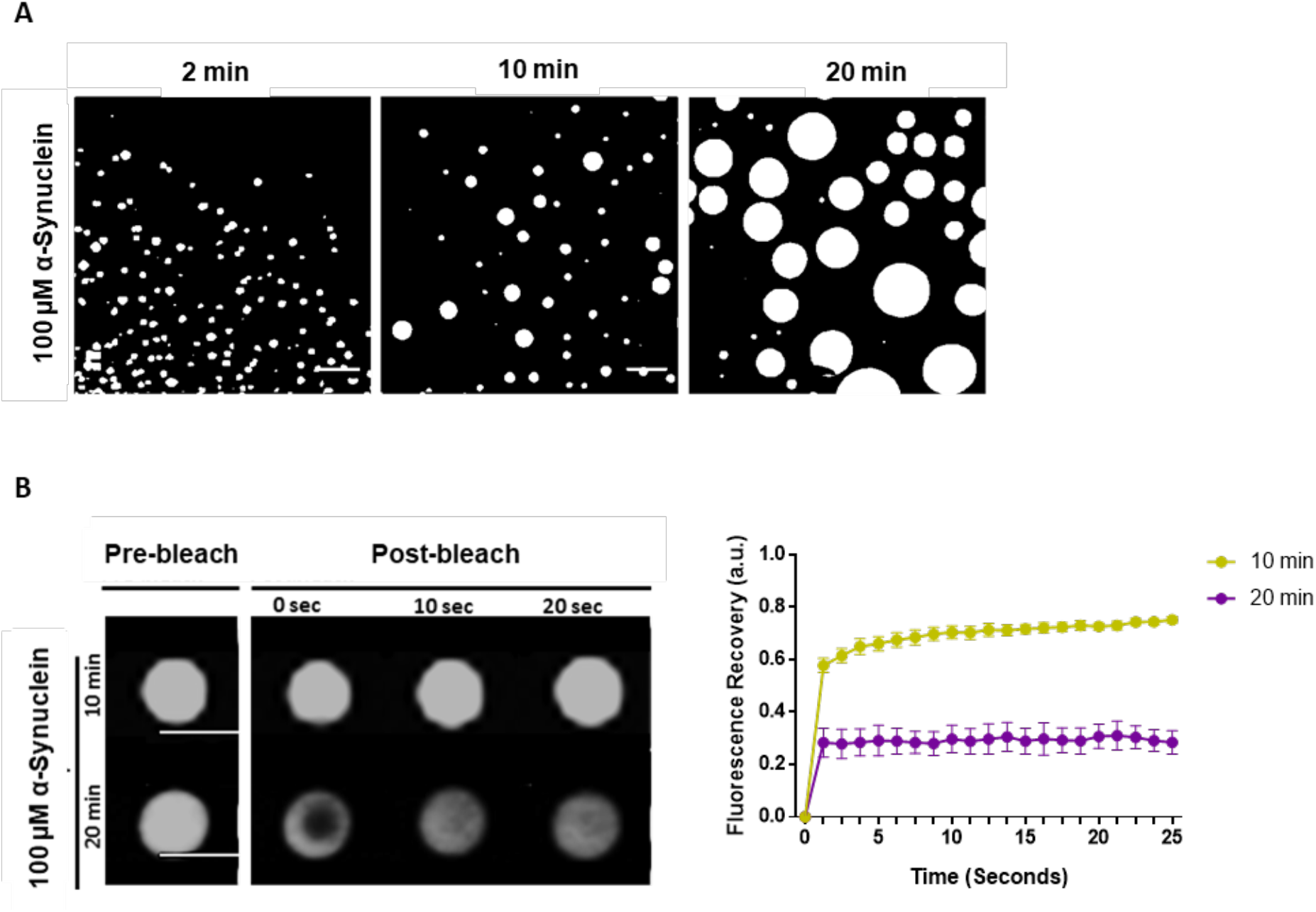
α-Synuclein condensates transitions from liquid to solid-like forms over time. (**A**) Fluorescence microscopy images displaying formation of α-synuclein condensates (100 μM) at 2, 10 and 20 min time points. Scale bar 10 μm. (**B**) FRAP measurements of 100 μM α-synuclein condensates at 10 and 20 min time points to measure the change in dynamics of condensates overtime. The images correspond to the region of interest with pre-bleach and post-bleach droplets at 0, 10 and 20 sec for each condition. Scale bar 5 μm. n= 4 per time range.

**Supplementary Figure 3.**
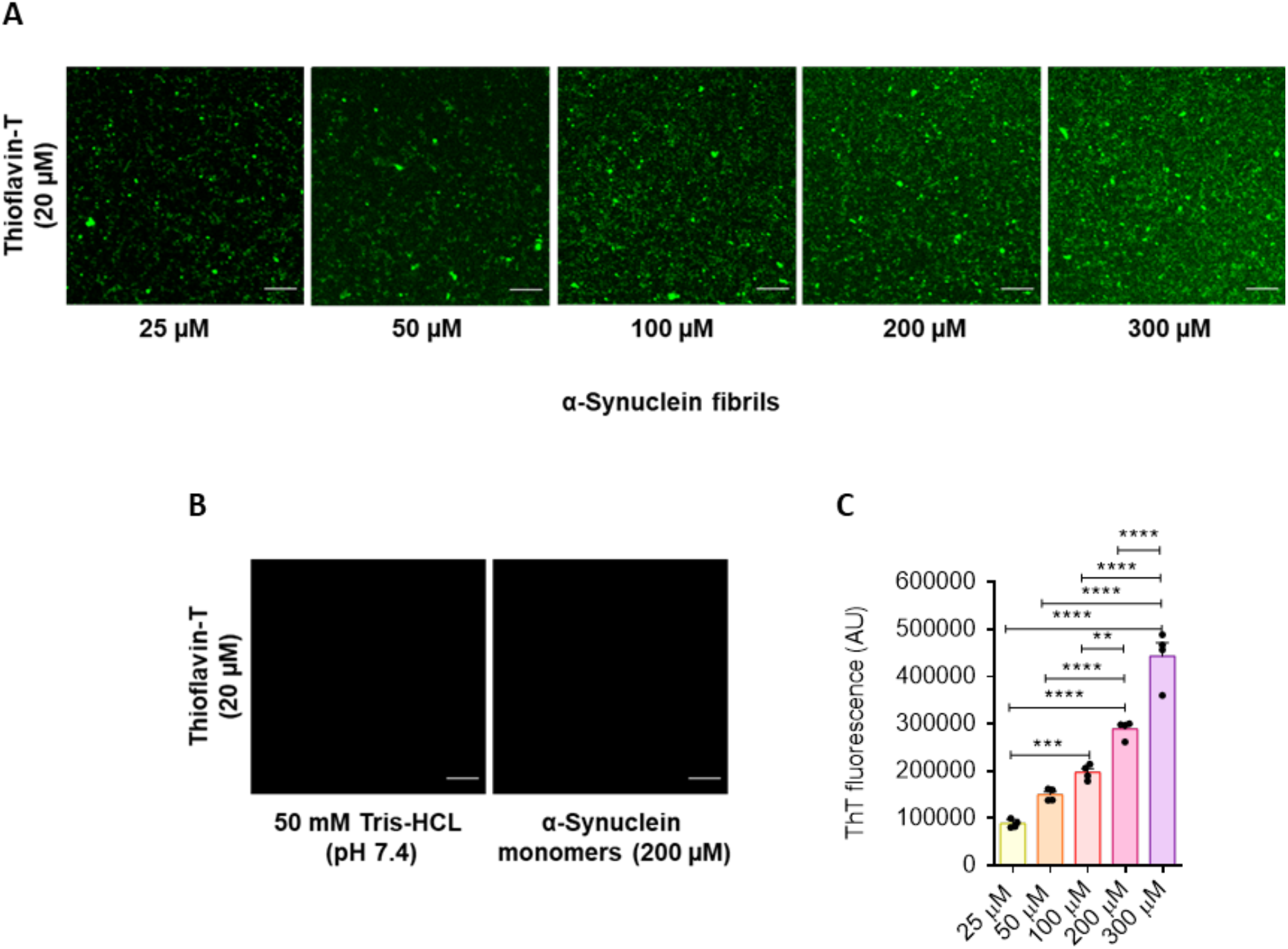
ThT can be used as a reporter dye for α-synuclein fibril formation with condensates. **(A)** Fluorescence microscopy images highlighting the concentration dependence of the ThT fluorescence intensity on the amyloid fibril load with increasing concentration of α-synuclein preformed fibrils (25, 50, 100, 200 and 300 μM) with 20 μM ThT in 50 mM Tris-HCl at pH 7.4. The scale bar represents 10 μm. **(B)** In the absence of amyloid structures, ThT fluorescence is not observed. Monomeric α-synuclein (200 μM) and 50 mM Tris-HCl at pH 7.4 in the presence of 20 μM ThT was used for control experiments. **(C)** Quantification of ThT (20 μM) amyloid-dependant fluorescence intensity of representative images shown in panel A for 25 μM (yellow), 50 μM (orange), 100 μM (red), 200 μM (magenta) and 300 μM (purple) α-synuclein preformed fibrils. All experiments were performed in 50 mM Tris-HCl (pH 7.4) in the presence 20 μM ThT. The data represent the mean ± SEM of n=4 individual experiments. One-way ANOVA., **P < 0.01, ***P < 0.001, ****P < 0.0001.

**Supplementary Figure 4.**
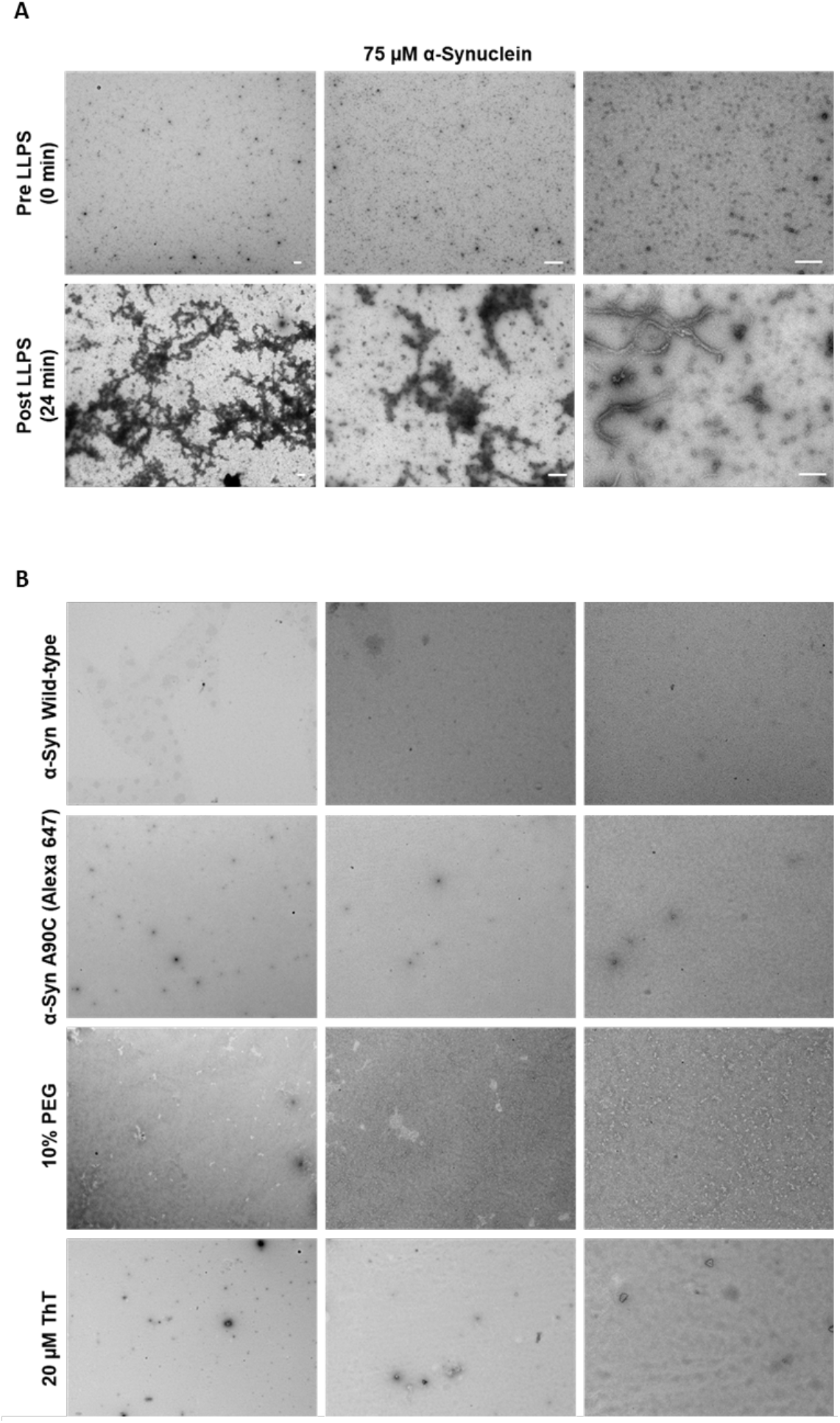
Transmission electron microscopy (TEM) shows that α-synuclein condensates evolve into aggregates over time and form cross-like network structures. **(A)** TEM images highlighting uranyl acetate α-synuclein species before (0 min) and after (24 min) liquid-liquid phase separation. Monomeric species are highlighted at 0 min, whilst fibrillar nature of α-synuclein post liquid-liquid phase separation are validated at 24 min. Experiments were performed with 75 μM α-synuclein in 50 mM Tris-HCl, pH 7.4, 10% PEG and 20 μM ThT. **(B)** TEM images showing the absence of aggregates after 24 mins of incubation with each individual component of the α-synuclein ThT aggregation assay through liquid-liquid phase separation. Experiments were performed in 50 mM Tris-HCl at pH 7.4. 1700x, 5000x and 11500x direct magnification from left to right panels were taken respectively. Scale bar represents 500 nm for all images.

**Supplementary Figure 5.**
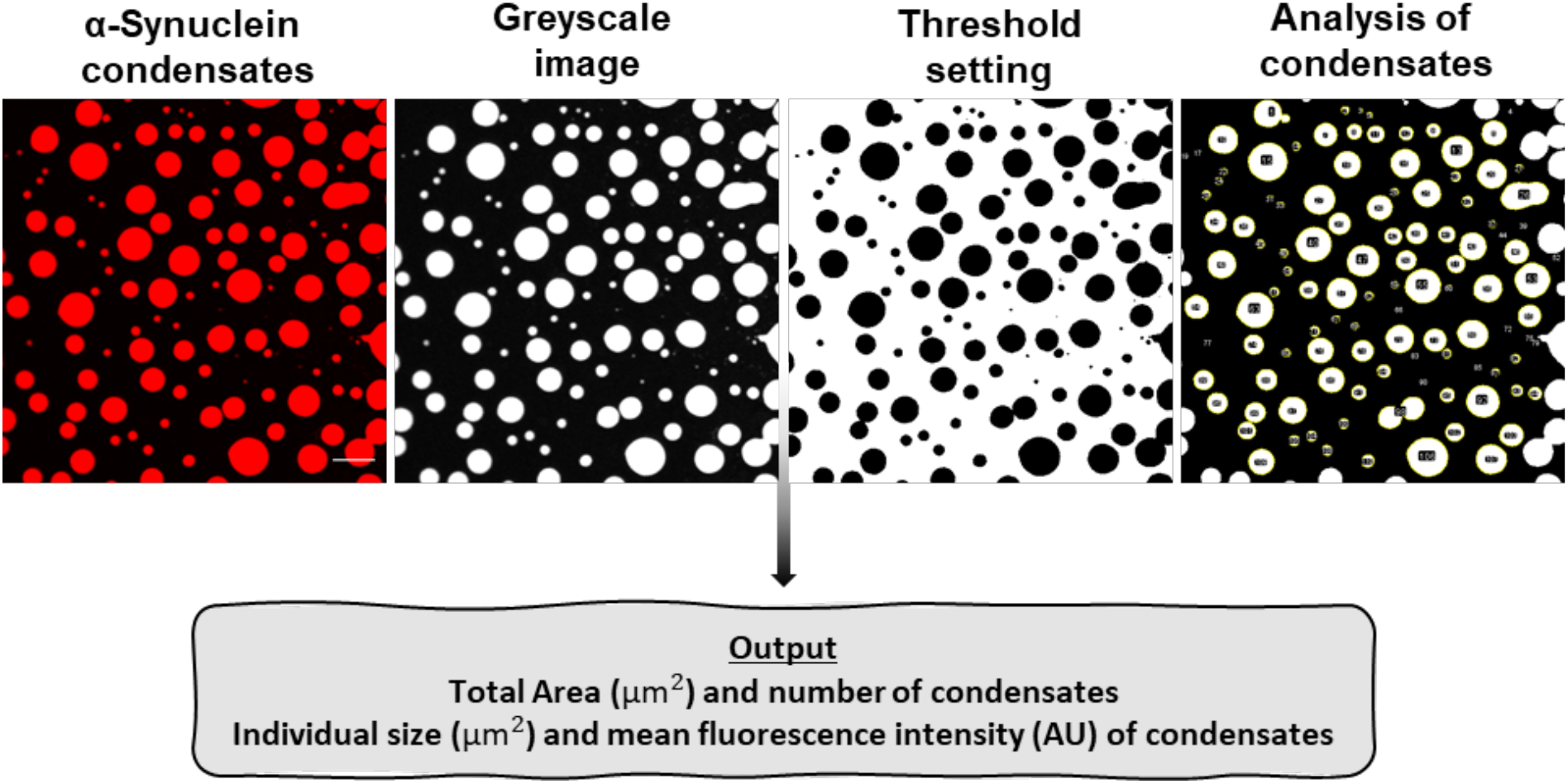
Fluorescence image analysis of α-synuclein condensates. Images of wildtype α-synuclein, containing 1 molar % A90C α-synuclein labelled with Alexa Fluor 647 were acquired on a Leica TCS SP8 inverted confocal microscope system. All images within an experiment were acquired using identical confocal settings (scan speed, resolution, magnification, laser intensity, gain, and offset). Images were analysed by applying threshold functions in ImageJ software that identified the phase separated α-synuclein condensates and excluded the background of the image. All condensates within the threshold limits were analysed for total area (μm^2^), number and average size of individual condensates (μm^2^) and mean fluorescence intensity of individual condensates (arbitrary units). The scale bar represents 10 μm.

**Supplementary Figure 6.**
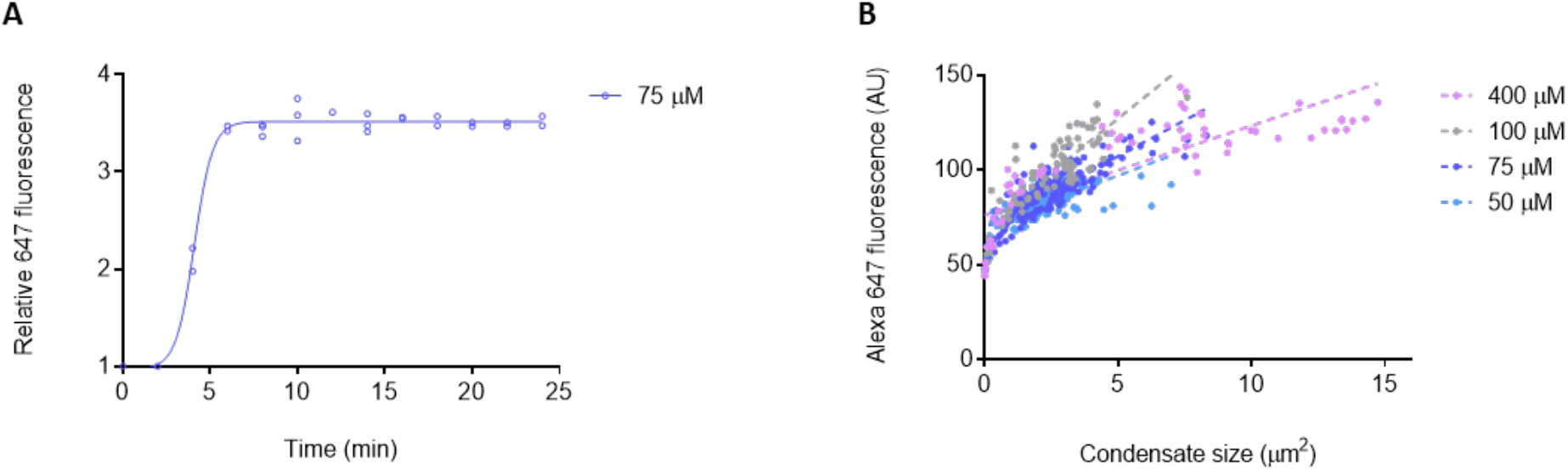
The Alexa Fluor 647 fluorescence varies only slightly within condensates. **(A)** Progression of Alexa Fluor 647 fluorescence intensity with time for α-synuclein (75 μM), which was measured at time points > 1 min and normalised against the first time point. After 5 min from the onset of liquid-liquid phase separation, the Alexa Fluor 647 fluorescence intensity remains constant. **(B)** Scatter plot showing the relationship between condensate size and Alexa Fluor 647 fluorescence for individual condensates of 400 μM (light purple), 100 μM (grey), 75 μM (blue) and 50 μM (cyan) α-synuclein, 10 min from the onset of liquid-liquid phase separation (n > 70 condensates per concentration); the linear regression coefficients r^2^ are 0.63 for 400 μM, 0.71 for 100 μM, 0.70 for 75 μM and 0.44 for 50 μM. Data are from a representative experiment that was repeated three times with similar results. All liquid-liquid phase separation experiments were performed in 50 mM Tris, pH 7.4 and 10% PEG.

**Supplementary Movie 1. Formation of α-synuclein condensates (75 μM) at the edge of a drop in 50 mM Tris, pH 7.4 and 10% PEG; the scale bar represents 20 μm.**

**Supplementary Movie 2. Fusion events of several α-synuclein condensates after 7 min post the onset of liquid-liquid phase separation; the scale bar represents 20 μm,**

**Supplementary Movie 3. Overlay of α-synuclein condensates formation and maturation, and α-synuclein aggregation at near-physiological concentration (75 μM) and conditions (50 mM Tris, pH 7.4) in the presence of 20 μM ThT; the scale bar represents 30 μm.**

